# Layer-specific transcriptomic changes across astrocyte subtypes in the aging human prefrontal cortex

**DOI:** 10.64898/2026.11.27.769633

**Authors:** Chih-Kai Chen, Chinh P Doan, Noor Sohail, Jane Banahan, Irais Valenzuela-Arzeta, Clemence Girardet, Shannan Ho Sui, Anupama Rai, Maria Mavrikaki

## Abstract

During aging, brain cell types including astrocytes undergo transcriptional changes associated with functional decline. As the most abundant glial cells in the brain, astrocytes form functional and metabolic connections with endothelial cells to maintain blood-brain barrier, provide metabolic and trophic support to neurons, and coordinate with microglia to clear of the cellular debris. This functional diversity is reflected in the transcriptional heterogeneity revealed by single-nucleus RNA sequencing across physiological and pathological conditions. However, it remains unclear how spatially distinct astrocyte subtypes age in different cortical microenvironments. Our recent high-resolution spatial transcriptomics analysis of the human dorsolateral prefrontal cortex (DLPFC) across the adult lifespan (Rai et al. 2026) identified more age-related differentially expressed genes (DEGs) in astrocytes than in any other glial cell types, along with age-related changes in astrocyte abundance. By sub-clustering two major astrocyte populations, in that study, we identified eleven functionally and spatially distinct astrocyte subtypes and characterized their layer-specific changes in proportion with aging. In this study, we further characterize these subtypes based on age-related differential gene expression profiles and enriched biological pathways. The most pronounced age-related response was observed across cortical layers 2-6 within astrocyte subtypes characterized as homeostatic, glia-associated, and stress-associated. Broadly, aging astrocytes demonstrated increased reactive features alongside decreased neuronal support programs. Pathways related to cellular stress and DNA damage were significantly upregulated, whereas actin-related pathways were significantly downregulated with aging. Together, these findings define astrocyte subtype- and layer-specific molecular programs associated with aging in the human DLPFC and identify candidate pathways that may contribute to age-related functional decline.

## Introduction

Higher cognitive functions in primates emerge from the dynamic interaction between neurons and glial cells. Age-related decline in working memory is closely associated with structural and functional changes in the dorsolateral prefrontal cortex (DLPFC) (Morrison and Baxter 2012), including changes in astrocytes that may contribute to altered neuronal support and circuit function. The DLPFC is organized into six cortical layers and the underlying white matter, each containing distinct neuronal and glial cell populations (Bayraktar et al. 2020). Understanding the spatially organized cellular and molecular mechanisms underlying age-related cognitive decline is critical for developing interventions that could help delay, prevent, or potentially reverse functional decline. However, the precise layer- and cell type-specific changes underlying its age-related functional decline, including the contribution of astrocyte heterogeneity, remains poorly understood.

Astrocytes are highly abundant glial cells in the brain and play a pivotal role in maintaining homeostasis, by supporting blood-brain barrier, providing metabolic and trophic support to neurons, and coordinating with microglia to clear cellular debris (Escartin et al. 2021; Hennes et al. 2025). Despite their functional diversity, astrocytes have traditionally been classified into two major morphological type: fibrous and protoplasmic astrocytes (Khakh and Deneen 2019). More recent studies have revealed the existence of distinct astrocyte subtypes within the rodent neocortex in morphology, transcriptomic profile, and function. Interestingly, their spatial distribution closely aligns with neuronal layers, potentially reflecting reciprocal neuron-astrocyte interaction (Batiuk et al. 2020; Bayraktar et al. 2020; Lanjakornsiripan et al. 2018). Astrocyte heterogeneity is even more pronounced in the human brain, with spatially-specific molecular profiles that likely reflect their diverse functions (Hodge et al. 2019; Man et al. 2024; Siletti et al. 2023).

Although the cerebral cortex is among the brain regions particularly vulnerable to aging and age-related neurodegenerative disease, many previous studies of aging astrocytes have relied on bulk-tissue analyses that lack laminar resolution (Boisvert et al. 2018; Sloan et al. 2026; Soreq et al. 2017). Thus, it remains unclear how distinct microenvironment across the cortical layers shape astrocyte subpopulations during aging.

In this study we build upon our recent work characterizing the cellular composition of each cortical layer and the underlying white matter in the human DLPFC across the adult lifespan using spatial transcriptomics (Rai et al. 2026). In that study, we showed that aging affects all cortical layers and white matter, with age-related transcriptomic changes reflecting both shifts in cell-type abundance and altered gene expression across multiple cell types, including astrocytes. We also identified eleven transcriptionally distinct astrocyte subtypes, each exhibiting a characteristic spatial distribution, with the abundance of several subtypes changing with aging. In this study, leveraging our high-definition spatial transcriptomics dataset (Rai et al. 2026) from 15 donors spanning 18-97 years of age, we further characterize astrocyte heterogeneity in the human DLPFC and investigate how these astrocyte subtypes change with aging across cortical layers, with a particular focus on layer- and astrocyte subtype-specific transcriptional remodeling.

## Results

### Functional characterization of spatially distinct astrocyte subtypes in the human DLPFC

We recently identified eleven spatially distinct astrocyte subtypes in postmortem human DLPFC tissue from fifteen individuals spanning the adult lifespan (18-97 years old) (Rai et al. 2026). To further characterize these spatially distinct astrocyte subtypes, we reanalyzed the expression of subtype marker genes identified in our previous study (Rai et al. 2026) separately in young (N=5, 18-40 years) and old (N=6, 70+ years) samples (**Figure 1A**), whereas the previous analysis examined these markers across the full cohort (N=15). We found that the marker genes for astro_1 to astro_10 were robustly expressed in both young and old brains, with greater subtype specificity observed in the older group. The greater subtype specificity of marker expression in older brains suggests that aging may promote more transcriptionally distinct astrocyte states, with reduced overlap among subtype-associated molecular programs. Astro_11 was identified in only two young brains, likely reflecting interindividual variation in tissue sampling and the specific cortical regions represented across specimens or rare transcriptional state only appearing in young individuals. We plotted average proportion of each subtype in each cortical layer in the young and old group (**Figure 1B**), and as noted in our earlier paper (Rai et al. 2026), we observed age specific changes in abundance in astrocyte subtypes in specific layers.

**Figure 1.**
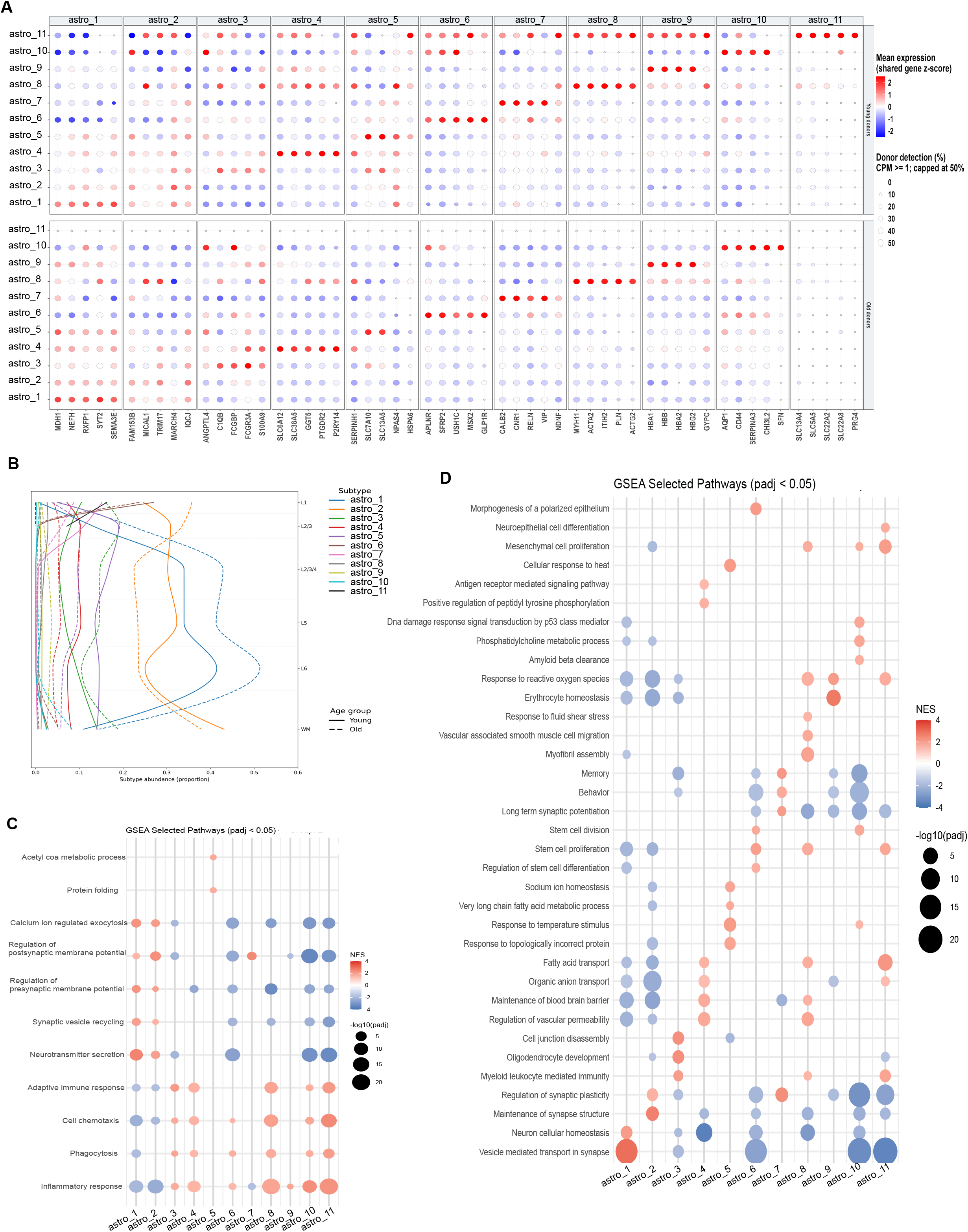
Characterization of 11 distinct astrocyte subtypes. **(A)** Expression of marker genes in all 11 astrocyte subtypes in young (upper panel) and old (lower panel) groups. **(B)** Spatial distribution of the average proportion of the 11 astrocyte subtypes across cortical layers in young and old samples. **(C)** Characterization of the 11 astrocyte subtypes using select significant GO:BP pathways related to homeostatic and reactive biological functions of astrocytes. **(D)** Dot plots showing select significant GO:BP pathways characterizing each of the 11 astrocyte subtypes based on enrichment of pathways related to homeostatic, stress-related, glia-associated, and vascular-associated, aging-related functions.

To functionally characterize these eleven astrocyte subtypes, we performed Gene Set Enrichment Analysis (GSEA) by using the subtype-specific marker genes ranked by their log fold change (**Figure 1C, D**). We first identified pathways associated with homeostatic and reactive astrocyte states and found that the eleven subtypes broadly segregated into two major functional groups, supporting their distinct subtype identities. Subtypes astro_1 and astro_2, and, to a lesser extent astro_7, were enriched for homeostatic pathways, while the other subtypes were enriched for pathways related to phagocytosis, and inflammatory response (**Figure 1C**). We identified that in our dataset the pro-inflammatory pathways were associated with genes including *CEBPB*, *LGALS3*, and *EDN1*, known regulators of astrocyte reactivity, gliosis, and apoptotic signaling (Gadea, Schinelli, and Gallo 2008; Wang et al. 2026). Well-known effectors of immune responses such as *C1R*, *CSF1*, and *CXCL16* were also present. The homeostatic pathways were characterized by genes encoding neuronal and synaptic proteins, such as *SYT3*, *VAMP2*, and *GABRA5*.

Next, we examined uniquely enriched pathways to further characterize each subtype (**Figure 1D**) and found that astro_1 and astro_2 were enriched for pathways related to maintenance of synapses and support for neurons. Astro_1 was characterized by enrichment of pathways related to vesicular transport including expression of genes encoding synaptotagmin and V-type ATPases, whereas astro_2 showed greater enrichment of structural maintenance programs, including genes such as *SYNGAP1* and junctophilin. Hereafter, we will refer to these subtypes as homeostatic type A and homeostatic type B astrocytes, respectively.

Astro_3 was characterized by pathways associated with interactions with different glial cell types (**Figure 1D**), particularly microglia and oligodendrocytes. This subtype expressed genes implicated in astrocyte-microglia crosstalk, including complement-related genes and *CX3CR1* (Matejuk and Ransohoff 2020), as well as genes associated with oligodendrocyte-related functions including *CLU* and *LYN* (Chen et al. 2024; Colognato et al. 2004). Hereafter, we refer to astro_3 as glia-associated astrocyte subtype.

Astro_4 was characterized by enrichment of stress-response pathways together with expression of genes associated with the blood-brain barrier (BBB) function (**Figure 1D**). This subtype expressed genes encoding members of the ABC and SLC transporter families, which are involved in transport processes at the BBB (Morris, Rodriguez-Cruz, and Felmlee 2017), as well as junction-associated genes including *CLDN5* and *CDH5* (Knox et al. 2022). Given the broader enrichment of stress-related pathways, we will refer to astro_4 hereafter as stress-associated astrocyte subtype with BBB-related features.

Astro_5 was characterized by expressing genes associated with stress response (**Figure 1D**), notably genes encoding heat shock protein family members including *SERPINH1* and *BAG3*, together with genes involved in ion homeostasis, such as *ATP1B2*. This subtype also expressed genes related to lipid metabolism, including *ABCD2* and *ACSBG1*, as well as genes previously associated with aging and reactive astrocyte states such as *ELOVL* family and *EDNRB* (Ferrero et al. 2025; Hammond et al. 2015). Hereafter, consistent with our earlier characterization (Rai et al. 2026) we will refer to astro_5 as stress-related astrocyte subtype.

Astro_6 was characterized by enrichment of pathways related to cell growth, proliferation, and differentiation, together with expression of genes involved in vascular signaling (**Figure 1D**). This subtype expresses numerous growth-related factors, including *SOX9*, *NOTCH1*, and *YAP1*. *SFRP2* is a soluble regulator of the Wnt and BMP pathways which has been reported to regulate stem cell functions (Alfaro et al. 2010). Consistent with its marker profile (Rai et al. 2026), we will refer to astro_6 hereafter as the vascular-associated astrocyte subtype A.

Astro_7 was characterized by enrichment of pathways related to higher cognitive functions, including long-term potentiation and memory (**Figure 1E**). This subtype also expressed several neuronal-associated genes, such as *RELN* and acetylcholine receptor genes. Previous single-cell RNA sequencing study in animal models has reported expression of neuronal genes in subsets of astrocytes (Schroeder et al. 2025), suggesting that astro_7 may represent a related transcriptional state. Hereafter, we refer to astro_7 as neuro-associated astrocyte subtype.

Astro_8 was characterized by enrichment of pathways associated with vascular smooth muscle cell (VSMC) function, including myofibril assembly and responses to fluid shear stress (**Figure 1D**). This subtype also expressed VSMC-associated genes, including *MYOCD* and *MYH11*. Hereafter, we refer to astro_8 as vascular-associated astrocytes subtype B.

Consistent with our previous marker gene-based characterization (Rai et al. 2026), Astro_9 was characterized by enrichment of erythrocyte-related pathways (**Figure 1D**) and expression of hemoglobin genes such as *HBA* and *HBB*. Hereafter, we will refer to astro_9 as the vascular-associated astrocyte subtype C.

Astro_10 was characterized by enrichment of aging-related pathways (**Figure 1D**), including clearance of amyloid-beta, downstream pathways of P53, and phosphatidylcholine metabolism. These were driven by genes well-known for being protective against pathological aging, including *TTPA* and *LRP2* (Nishida et al. 2009; Sagare et al. 2007), and genes involved in lipid metabolism, such as *SLC44A3* and *LCAT*. Hereafter, we refer to astro_10 as aging-reactive astrocyte subtype.

Astro_11, as previously reported (Rai et al. 2026), was a rare subtype represented by a very limited number of bins and was spatially restricted to the upper cortical layers. It was characterized by pathways related to vascular remodeling, mesenchymal cell proliferation and neuroepithelial cell differentiation (**Figure 1D**). Marker genes included matrix-remodeling and vascular-associated genes such as members of *MMP* family, *SERPINE1*, and SLC family transporters, and more upstream regulators such as *FGF10*, *FOXC2*, and *CCND1*. Consistent with its superficial distribution and our previous classification, we refer to astro_11 hereafter as astrocytes with progenitor cell-like transcriptional signatures.

### Cortical layer- and subtype-specific aging-related gene expression changes in human DLPFC astrocytes

We next investigated age-related layer- and subtype-specific changes in astrocyte gene expression. To address this, we performed differential gene expression (DGE) analysis for each subtype-layer combination by comparing old versus young samples.

We observed that aging-related transcriptional changes were primarily concentrated in astrocyte subtypes classified as homeostatic (types A and B), glia-associated, and stress-associated, which exhibited the largest numbers of significant differentially expressed genes (DEGs) across layers 2-6 (**Figure 2A**). In contrast, astrocyte subtypes in layer 1 and white matter showed relatively few aging-related gene expression changes. Several subtype-layer combinations, including certain aging-reactive and homeostatic type A astrocytes in layer 1, did not meet the minimum threshold required for DEG analysis (N ≥ 50 bine per sample in ≥ 3 samples) and are therefore not shown in Fig 2A. We further focused on astro_1 to astro_5 which showed the most number of DEGs across the cortical layer 1-6 and white matter. We next plotted log fold change against the padj (**Figure 2B**, **2C**) to analyze effect size and expression trends. This analysis revealed global upregulation of *GFAP* and a widespread downregulation of *SEMA6B* and *TNR* across the subtypes astro_1, astro_2, astro_3, astro_4 astro_5 (**Figure 2B**, **2C**) in L2/3 and L2/3/4. We also identified unique gene signatures for the subtypes with most DEGs within cortical layer 2/3 and L2/3/4 (**Figure 2D**, **2E**).

**Figure 2.**
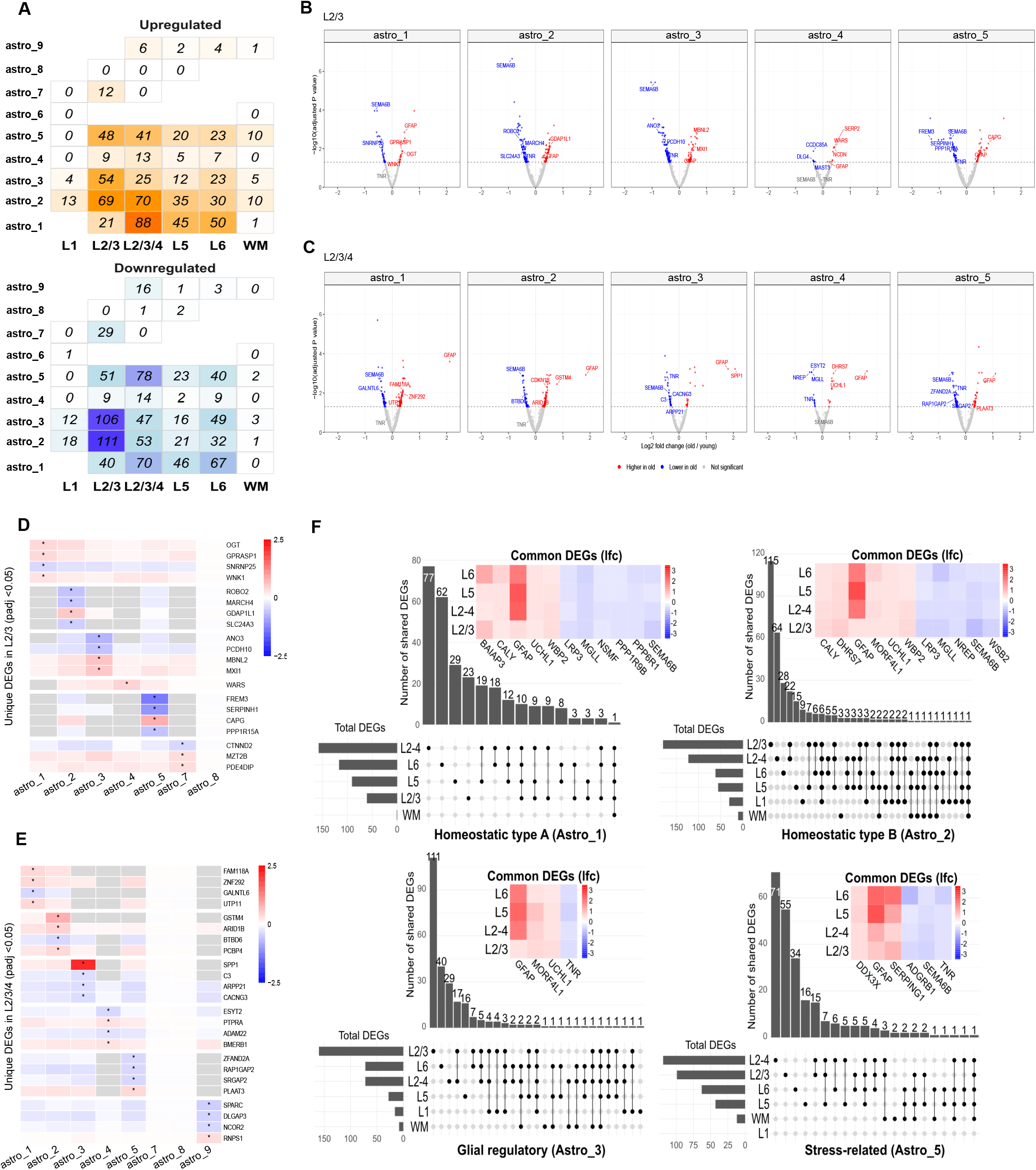
Aging-related differentially expressed genes (DEGs) across astrocyte subtypes. **(A)** Number of significant DEGs in each subtype-layer. Subtype-layer combinations with less than 3 young or old samples having at least 50 bins were excluded from the analysis. (**B, C**) Volcano plot showing DEGs in L2/3 and L2/3/4. Select unique and widely dysregulated genes across astrocyte subtypes are highlighted. **(D, E)** Astrocyte subtype specific unique genes in L2/3 (D) and in L2/3/4 (E). **(F)** Upset plots of significant DEGs in the four selected subtypes. Heatmaps reflect the log2 fold change (lfc) values of genes shared across layers 2-6 within each subtype.

To identify shared and layer-specific DEGs across cortical layers within individual astrocyte subtypes, we performed an intersection-overlap analyses (**Figure 2F**). We found that, for most astrocyte subtypes, a larger proportion of aging-related DEGs was unique to individual cortical layers than shared across multiple layers (**Figure 2F**), indicating that the transcriptional response of a given astrocyte subtype to aging is strongly influenced by its laminar context.

Although many aging-related DEGs were unique to specific astrocyte subtypes, they often converged on similar biological processes. For example, in cortical layers 2-4, homeostatic type A (astro_1) and B (astro_2) and glia-associated (astro_3) astrocytes showed upregulation of genes involved in cellular stress responses and neuroprotective pathways associated with neurodegeneration. *OGT*, a suppressor of astrocyte reactivity, and *SPP1*, a secreted inhibitor of neurodegeneration (Dong et al. 2023; Li and Jakobs 2022), were upregulated in these layers. *MXI1*, a downstream effector of HIF signaling, which is involved in responses to hypoxic stress (Pantazopoulou et al. 2021), was also increased, together with other stress-related genes such as *GSTM4*.

In contrast, layers 5 and 6 showed downregulation of the protein homeostasis-related gene *DNAJB5* (Zarouchlioti et al. 2018). The stress-associated astrocyte subtype also exhibited downregulation of genes involved in cellular stress responses, including *PPP1R15A* and *ZFAND2A* (Lee, Takayama, and Goldberg 2018; Smith et al. 2020), as well as downregulation of the senescence-related gene *ANGPTL2* in layer 6 (Thorin-Trescases et al. 2021). Together, these findings suggest substantial layer- and subtype-specific transcriptional remodeling in astrocytes during aging.

### Layer- and subtype-specific enrichment of homeostatic and stress-related pathways with aging in human DLPFC astrocytes

To identify biological pathways associated with aging-related DEGs, we performed GSEA of each subtype-layer combination. The highest numbers of significantly altered pathways were observed in homeostatic types A and B, glia-associated and stress-associated subtypes within cortical layers 2/3, 2/3/4, 5, and 6. In layers 2/3, pathways related to cellular stress responses, including autophagy, oxidative stress, and metal ion homeostasis, were predominantly upregulated with aging (**Figure 3**), particularly in glia-associated (astro_3) and stress-associated (astro_5) astrocyte subtypes.

**Figure 3.**
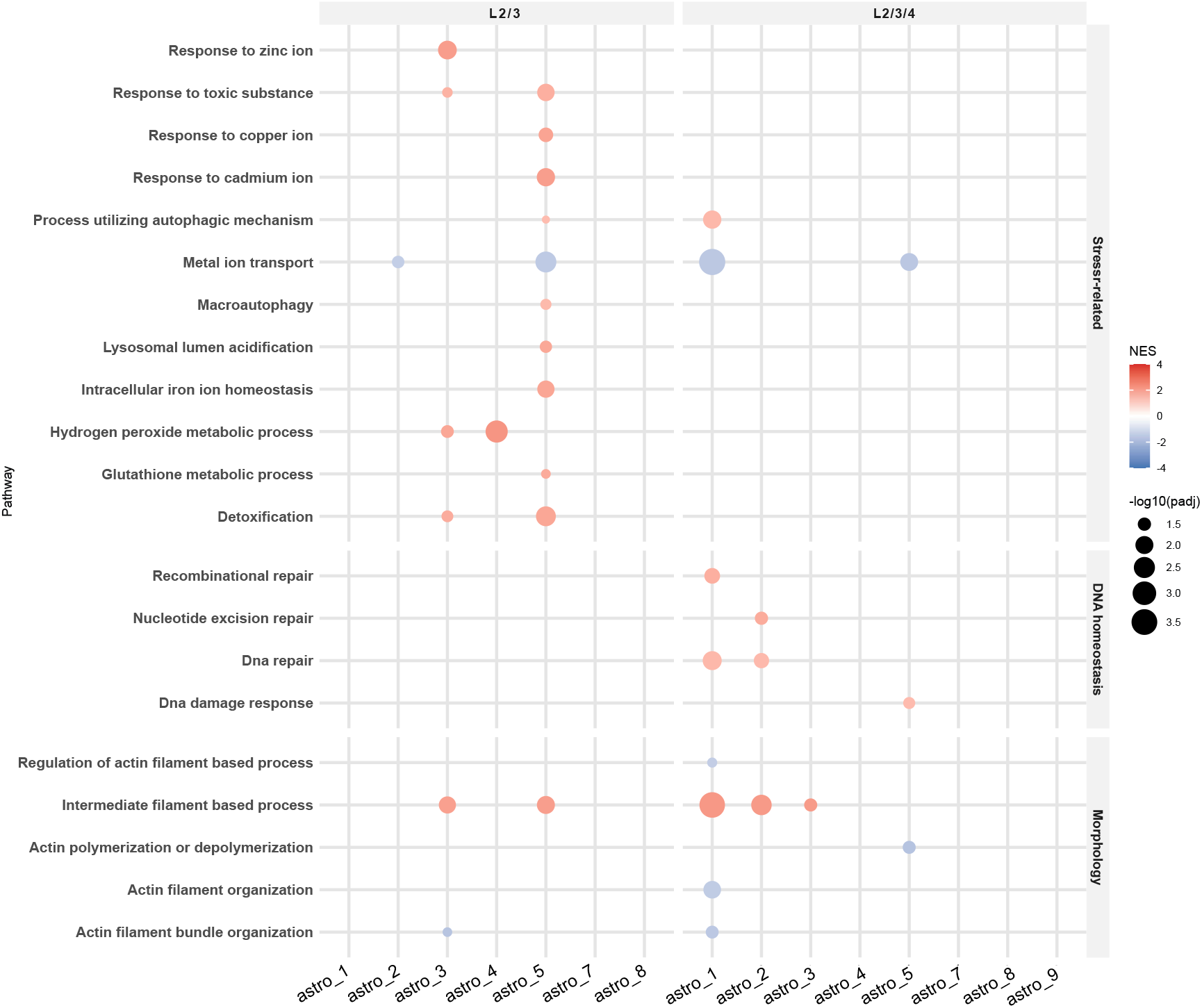
Aging-related Gene Ontology (GO) analyses across astrocyte subtypes. Dot plots of selected significantly changing GO:BP pathways in selected cortical layers and astrocyte subtypes. Pathways are grouped by their functions.

We also observed a coordinated upregulation of pathways related to DNA homeostasis. Pathways involved in DNA repair, such as nucleotide excision repair and double strand break repair, tended to be upregulated in layers 2/3/4. Notably, these DNA damage-related pathways were not significantly altered in layer 2/3, where stress-related pathways predominated.

We next examined morphology-related pathways, given previous reports of age-associated changes in astrocyte morphology. Driven in part by upregulation of *GFAP* expression, the “intermediate filament-based process” pathway was upregulated across most subtype-layer combinations, whereas actin-related pathways were downregulated. Overall, GSEA revealed that stress-response and DNA repair programs were engaged in a layer- and subtype-restricted manner, whereas cytoskeletal remodeling was a near-universal feature of astrocyte aging.

## Discussion

Recent advancements in high-throughput transcriptomic approaches, including single-cell and single-nucleus RNA sequencing, have revealed substantial regional transcriptional heterogeneity among astrocyte populations. However, because astrocytes are shaped by the local brain microenvironment, these sequencing approaches fail to preserve the spatial context needed to resolve spatially and transcriptionally distinct astrocyte populations within a given region. In this study, we leveraged high-resolution spatial transcriptomic data from the human DLPFC to further characterize the functional profiles of spatially distinct astrocyte subtypes identified in our recent work (Rai et al. 2026). We classified these eleven spatially and transcriptionally distinct astrocyte subtypes into functional categories, including homeostatic (types A and B), glia-associated, stress-associated, barrier-associated, aging-associated, and vascular-associated subtypes based on the enrichment of pathways linked to established astrocyte functions. Recent studies using single-nucleus RNA sequencing data from aged and Alzheimer’s disease cohorts have similarly partitioned astrocytes into 9-10 (Green et al. 2024; Yang et al. 2024) subpopulations, categorizing them as homeostatic, reactive, intermediate, stress, interferon-responsive, among others (Green et al. 2024; Yang et al. 2024). However, because these studies relied on dissociated nuclei from aged and AD populations, they lacked both the cytoplasmic transcripts captured by our spatial transcriptomic approach and representation across younger and middle-adult ages. Thus, our study extends prior work by defining spatially distinct astrocyte transcriptional subtypes across the full adult lifespan.

Astrocyte subtypes characterized as homeostatic (types A and B), stress-associated, and neuron-associated were predominantly localized to the cortical gray matter (layers L2-L6) (**Figure 1B**), whereas vascular subtypes B and C exhibited more spatially clustered distributions, consistent with their distinct functional profiles. These findings support the functional relevance of astrocyte transcriptomic heterogeneity and demonstrate that distinct transcriptional states are associated with discrete spatial distributions within the human cortex.

Among the eleven subtypes, aging-associated astrocytes were enriched for pathways related to amyloid clearance and lipid metabolism, particularly phosphatidylcholine (PC) metabolism. PC has been implicated in mitochondrial aging in *C. elegans* models and reduced circulating PC levels have also been reported in Alzheimer’s disease patients (Poliezhaieva et al. 2026; Whiley et al. 2014). It has also been reported that astrocyte PC metabolism could be involved in dendrite maintenance and astrocyte reactivity (Xu et al. 2016; Zhu et al. 2016). While the exact role of PC in aging is yet obscure, dysregulated lipid metabolism in astrocytes is a well-established hallmark of neuroinflammation and senescence (Gildea and Liddelow 2025).

We also found that homeostatic (type A and B), glial-associated, and stress-associated astrocyte subtypes were the most affected by aging, particularly across the cortical layers 2-6. Notably, our DE analyses were restricted to subtype-layer combinations represented by at least three young and three old samples, with minimum of 50 bins per sample, thereby reducing the likelihood that observed age-related differences were driven by sparsely represented populations. Consequently, certain rarer groups—such as the aging-associated and meningeal connective tissue subtypes—were excluded from this DE analysis as they didn’t meet those criteria. Therefore, future studies utilizing larger datasets will be essential to validate these subtypes and explore their relationship with pathological aging.

Among the selected subtypes and layers, we noticed a global increase in *GFAP* and decrease in *SEMA6B* and *TNR*. *GFAP* is a marker for reactive astrocytes, and its increase is a well-known reaction of astrocytes during aging (Nichols et al. 1993); while *SEMA6B* and *TNR* are involved in neuronal support functions such as axonal growth or myelination (Cordovado et al. 2022; Wagner et al. 2020). Genes related to protein and RNA homeostasis have also been identified among the most robustly changed genes in many subtype-layers. These findings suggest increased reactivity and reduced homeostatic capacity to be among the most shared aging responses across all astrocyte subtypes and cortical layers.

We further grouped GSEA identified pathways into stress response-related pathways, DNA damage response-related pathways, and morphology-related pathways as aging-related processes. These pathways also exhibited enrichment in specific subtypes or layers during aging. For instance, cellular stress responses, particularly oxidative stress and proteostasis-related pathways, are well-established hallmarks of senescent astrocytes (Cohen and Torres 2019; Gildea and Liddelow 2025). Senescence in astrocytes has been associated with defective autophagy and results in a specific subtype of autophagy-dysregulated astrocytes (APDAs) under disease conditions (Cabral-Miranda, Matias, and Gomes 2025; Lee et al. 2022). In our dataset, however, we saw an upregulation of autophagy pathways along with these stress response pathways in glia-associated and stress-associated subtypes (**Figure. 3**), suggesting differential regulation of autophagy-related pathways during physiological versus pathological aging. Furthermore, metal ions, such as iron, zinc, and copper, are known modulators of brain aging and neurodegeneration (Chen et al. 2025). The levels of these metal-ions is known to be regulated by astrocytes (James et al. 2012; Waller et al. 2018; Xu et al. 2017) and their imbalance leads to cellular stress responses, such as oxidative stress. While these pathways were predominantly enriched in L2/L3, our previous study also identified a significant reduction in excitatory L2/L3 neurons (Rai et al. 2026). Whether the upregulation of stress-related pathways in astrocytes contribute to neuronal loss or instead arise as a consequence of neuronal vulnerability remains unknown and will require further mechanistic investigation.

DNA damage is a well-established feature of brain aging and cellular senescence, including in astrocytes (Gildea and Liddelow 2025). Downstream to DNA damage, astrocytic P53 has been shown to regulate astrocytic senescence and neuronal apoptosis (Turnquist et al. 2016). We observed upregulation of pathways related to DNA homeostasis such as DNA repair predominantly in homeostatic astrocyte subtype in layers 2-4. Because astrocytes play a central role in brain energy metabolism, which can generate reactive oxygen species, they may be particularly vulnerable to age-related oxidative stress and DNA damage.

Astrocytes in the aged brain are known to upregulate genes related to complement pathway such as *C3*, *C4b*, and *C1q* (Clarke et al. 2018; Palmer and Ousman 2018). While in our dataset upregulation of these complement genes during aging was not observed, we saw upregulation in other hallmark genes such as *SERPINA3* and *GFAP*. It is possible that by subdividing astrocytes into 11 transcriptionally distinct subtypes distributed the broader age-related increase in reactivity across multiple populations, thereby reducing the effect size and statistical power within individual subtype comparisons. On the other hand, the increases in astrocytic complement-related genes reported in previous studies may be more pronounced in other brain regions than in the cortex (Boisvert et al. 2018; Clarke et al. 2018). In addition, most astrocyte subtypes displayed impairment of neuron-supportive pathways together with changes in genes involved in proteostasis and RNA modification, features that are also associated with senescence in astrocytes (Cohen and Torres 2019; Gildea and Liddelow 2025). These findings raise the possibility that, despite a broader in pro-inflammatory gene expression, astrocytes in the aging cortex may preferentially shift towards senescent-like rather than a predominantly reactive phenotype during physiological aging.

Astrocyte morphology is related to its function (Baldwin, Murai, and Khakh 2024), and changes in astrocyte morphology have been reported in aging and in disease states. Additionally, *GFAP*, whose levels have been shown to increase in reactive astrocytes, is an intermediate filament protein regulating astrocyte shape (Yang and Wang 2015). We observed upregulation of intermediate filament processes in all the layers and most of the astrocyte subtypes. However, we observed downregulation of pathways related to actin filament. While the exact functional link between astrocyte cytoskeletons, especially actin, with age-related processes is still obscure, actin microfilament has already been shown to be related to pro-inflammatory and age-related pathways (Villablanca, Vidal, and Gonzalez-Billault 2023). The Wnt signaling pathway is a known regulator of astrocyte morphology, notably via actin-related pathways, and affects astrocyte crosstalk with other cell types, including neurons and microglia (Faust et al. 2025; Szewczyk et al. 2024). While the role of Wnt signaling in aging is not yet well understood, other recent studies have also shown this pathway being affected in aging cortical astrocytes (Hennes et al. 2026).

Several methodological limitations of this study should be considered. First, our data were analyzed at a bin size of 16 μm, which approaches single resolution but may still capture a mixture of cells. Additionally, astrocytes have highly branched processes that can extend hundreds of microns and high-definition spatial transcriptomics using contiguous bins can capture transcripts localized not only in the soma but also throughout the cytoplasm and cellular processes, thus mapping more bins to astrocyte.as compared to nuclei based approaches Secondly, although brains from donors with no known neurological or psychiatric conditions were selected for this study, our inclusion criteria did not exclude individuals with peripheral medical conditions.

In conclusion, our study identifies spatially and functionally distinct astrocyte subtypes in the human DLPFC using high-resolution spatial transcriptomics. These analyses reveal broader age-related trends towards increased reactivity and decreased neuronal support, while also uncovering layer- or subtype-specific processes, including differential stress response programs. Future studies should define how these astrocyte subtypes interact with neighboring neuronal and glial populations and establish how they are altered during pathological aging. Functional validation in complementary *in vitro* and *in vivo* models will be important to determine the relevance of these transcriptional states. Together, these studies will help determine the contribution of astrocyte heterogeneity to human brain aging. Our study is significant because it advances our understanding of astrocyte heterogeneity in the human brain and highlights subtype-specific molecular pathways that may offer new therapeutic opportunities to mitigate age-related cognitive decline.

## Materials and Methods

### Data

We reanalyzed spatial transcriptomics data from fifteen DLPFC samples spanning the adult lifespan (Rai et al. 2026). All data were generated using the 10x Genomics High-Definition Visium (Visium HD) platform and analyzed at 16 μm resolution. The fifteen samples included five young (18-39 years old), four middle-aged (40-69 years old), and six old (70-97 years old) donors. Samples of all three age groups were previously included in the analysis that identified the eleven astrocyte subtypes (Rai et al. 2026), while only the young and old samples were included for aging-related DGE and GSEA analyses. Detailed methodology regarding data generation and processing can be found in our publication (Rai et al. 2026).

### Bioinformatics analysis

#### Astrocyte subtype marker gene expression

Expression of astrocyte subtype markers identified in our previous study (Rai et al. 2026) were plotted in the young and old samples using the pseudobulk data file.

### DGE analysis across the eleven astrocyte subtypes

To identify layer specific aging-regulated genes in astrocyte subtypes, we applied a pseudobulk approach. To aggregate the counts on the level of sample x age group x cortical layer x astrocyte-subtype, we used the AggregateExpression() function from Seurat. To be included in the analysis, we required that a subtype in a cortical layer have a minimum of 50 bins in at least 3 samples. We used the Bioconductor R package DESeq2 (Love, Huber, and Anders 2014) to test for DE of the counts, modeling the data using a combined variable called age_layer _astrocyte_subtype. After the model is fit, we apply shrinkage (“ashr”) so that gene log fold change (LFC) is not dependent overall on basal gene expression level.

### Average proportion of astrocyte subtypes in different cortical layers

The average proportion of each subtype per layer was calculated in all young or old samples. The thickness of each layer in the corresponding plot (**Figure 1B**) also corresponds to the average number of total bins of that layer. The abundance of each subtype was marked as dots at the vertical center of each layer and connected with smoothed lines using PchipInterpolator from the module scipy.interpolate in Python.

### Pathway enrichment analysis

We performed GSEA by ranking DEGs by log fold change and compared with the collection C5:GO:BP. For characterization of the eleven astrocyte subtypes, GSEA was performed on the marker gene list for each astrocyte subtype. For identifying changes in aging, GSEA was performed on the DEG list of each astrocyte subtype in each cortical layer, comparing young versus old. We then manually selected the relevant pathways shown in this article.

## Author Contributions

MM and AR conceptualized, designed, and supervised the study. Visium HD data reanalyzed in this study was generated by AR. CKC analyzed the data, generated the figures, and wrote the manuscript. NS, CPD, and IVA performed bioinformatic analysis of the data presented. JB assisted with tissue preparation and quality control assessment. CG contributed to the discussion of astrocyte biology. SHS supervised the bioinformatics analysis of the study. All authors reviewed, edited, and approved of the final manuscript.

The content is solely the responsibility of the authors and does not necessarily represent the official views of Harvard Catalyst, Harvard University and its affiliated academic healthcare centers, or the National Institutes of Health.

